# Unravelling Rubber Tree Growth by Integrating GWAS and Biological Network-Based Approaches

**DOI:** 10.1101/2021.08.16.456528

**Authors:** Felipe Roberto Francisco, Alexandre Hild Aono, Carla Cristina da Silva, Paulo de Souza Gonçalves, Erivaldo José Scaloppi, Vincent Le Guen, Roberto Fritsche Neto, Livia Moura Souza, Anete Pereira de Souza

**Author notes:** These authors have contributed equally to this work (Felipe Roberto Francisco, Alexandre Hild Aono). Correspondence: Anete Pereira de Souza.

## Abstract

*Hevea brasiliensis* (rubber tree) is a large tree species of the Euphorbiaceae family with inestimable economic importance. Rubber tree breeding programs currently aim to improve growth and production, and the use of early genotype selection technologies can accelerate such processes, mainly with the incorporation of genomic tools, such as marker-assisted selection (MAS). However, few quantitative trait loci (QTLs) have been used successfully in MAS for complex characteristics. Recent research shows the efficiency of genome-wide association studies (GWAS) for locating QTL regions in different populations. In this way, the integration of GWAS, RNA-sequencing (RNA-Seq) methodologies, coexpression networks and enzyme networks can provide a better understanding of the molecular relationships involved in the definition of the phenotypes of interest, supplying research support for the development of appropriate genomic based strategies for breeding. In this context, this work presents the potential of using combined multiomics to decipher the mechanisms of genotype and phenotype associations involved in the growth of rubber trees. Using GWAS from a genotyping-by-sequencing (GBS) *Hevea* population, we were able to identify molecular markers in QTL regions with a main effect on rubber tree plant growth under constant water stress. The underlying genes were evaluated and incorporated into a gene coexpression network modelled with an assembled RNA-Seq-based transcriptome of the species, where novel gene relationships were estimated and evaluated through in silico methodologies, including an estimated enzymatic network. From all these analyses, we were able to estimate not only the main genes involved in defining the phenotype but also the interactions between a core of genes related to rubber tree growth at the transcriptional and translational levels. This work was the first to integrate multiomics analysis into the in-depth investigation of rubber tree plant growth, producing useful data for future genetic studies in the species and enhancing the efficiency of the species improvement programs.

## **1** Introduction

*Hevea brasiliensis* (rubber tree) is an outbreeding forest species belonging to the Euphorbiaceae family with an inestimable importance in the world economy because it is the only crop capable of producing natural rubber with quantity and quality levels able to meet global demand (De Faÿ and Jacob, 1989). Possessing unique characteristics such as resistance, elasticity and heat dissipation, Hevea rubber is used as a feedstock for more than 40,000 products (Pootakham et al., 2017; Mantello et al., 2019). Although it is very important, *H. brasiliensis* is still in an early domestication stage due to its long breeding cycle (25 to 30 years), the large areas required for planting and its recent cultivation (Priyadarshan and Clément-Demange, 2004; Gonçalves et al., 2006). In this context, Hevea breeding programs aim to improve important agronomic traits for rubber fabrication, mainly those related to latex growth and production (Priyadarshan, 2003). The use of early genotype selection technologies has been proposed as a breeding alternative for accelerating this process, e.g., incorporating genomic tools for marker-assisted selection (MAS) (Pootakham et al., 2017; Priyadarshan, 2017). Although the discovery of quantitative trait loci (QTLs) can benefit Hevea breeding programs (Souza et al., 2019), this characterization is hindered by the large number of genes and molecular interactions controlling such characteristics (Pootakham et al., 2020). To date, few QTLs have been successfully used for rubber tree MAS for complex quantitative traits due to the insufficient quantity of linked markers in the QTLs, small QTL effects on the phenotype, or strong environmental influences (Nguyen et al., 2019).

Several studies have been carried out in the last decade to identify QTLs in *H. brasiliensis* through genetic linkage maps (Souza et al., 2013; Pootakham et al., 2015; Conson et al., 2018; Rosa et al., 2018; Xia et al., 2018) and association mapping (Chanroj et al., 2017). Genome-wide association studies (GWAS) are important tools for the identification of candidate genetic variants underlying QTLs, with great potential to be incorporated into MAS. Compared to linkage maps, the use of GWAS methodologies has advantages such as using genetically diverse populations with different rates of recombination and linkage disequilibrium (LD) (Myles et al., 2009). Despite the observed GWAS efficiency in several crops (Warraich et al., 2020; Zhang et al., 2020; Verzegnazzi et al., 2021), this methodology still presents limitations related to the low proportion of phenotypic variance explained by the identified genomic regions (Manolio et al., 2009). As an alternative, the combination of GWAS results with other molecular methodologies, such as transcriptomics and proteomics analyses, can contribute to better knowledge of the genetic mechanisms involved in the definition of a trait (Tam et al., 2019), overcoming the statistical limitations on the characterization of a broad set of causal genomic regions.

Although the identification of genes with a great phenotypic effect is consolidated with GWAS methodologies (Nebel et al., 2011), there are no established methods for investigating the complete set of genes controlling complex traits through multiomics approaches, and such characterization is an open scientific challenge, especially in crops with complex genomes such as rubber trees (Schaefer et al., 2018). Different initiatives have associated GWAS results with RNA-Seq data (Yan et al., 2020), linking causal genes relevant to the observed phenotypic variation with cell transcription activity profiles (Schaefer et al., 2018; Nguyen et al., 2019). In Hevea, however, RNA-Seq-based studies have been mainly performed to investigate differentially expressed genes (DEGs) under different environmental or stress conditions and profiling rubber tree samples (Hurtado Páez et al., 2015; Sathik et al., 2018; Mantello et al., 2019; Ding et al., 2020; Ding et al., 2020). Although the integration of GWAS with RNA-Seq methodologies has proven to provide a deeper comprehension of the genetic relationships involved in trait definition, there is no study, to date, aggregating such data in Hevea.

We are currently undergoing a major revolution in omics sciences (genomics, transcriptomics, proteomics and phenomics) with different methods for data integration enabling important advances in all phases of genetic improvement, ranging from the discovery of new variants to the understanding of important metabolic pathways (Scossa et al., 2021). The integration of data derived from multiomics can be combined to reveal, in a profound way, the relationships that represent the true biological meaning of the studied elements (Jamil et al., 2020; Wu et al., 2020). This approach has become increasingly common in humans (Wu et al., 2018), animals (Fonseca et al., 2018), microorganisms (Wang et al., 2019), and combinations of species (Pinu et al., 2019). However, for plants, such integrated methodologies are still a great challenge, especially for nonmodel species with elevated genetic diversity and complex genomes (Jamil et al., 2020), which is the case for *H. brasiliensis* (Tang et al., 2016; Liu et al., 2020, Wu et al., 2020). Despite its economic importance, no study incorporating multiomics has been carried out on *H. brasiliensis*. With the wide availability of omics data, coexpression networks have become a tool with great potential for inferring gene interactions, mainly based on regulatory and structural relationships, allowing for a broader understanding of unknown molecular mechanisms (Rao and Dixon, 2019). The identification of these genes also allows us to indirectly assess, through their enzymes, the global metabolic relationships involved in defining the evaluated characteristic (Pérez-Bercoff et al., 2011). In this way, we can make use of GWAS to select genes of great importance for the phenotype of interest. Such genes can be used as a guide to select modules of coexpressed genes and their enzymes, which may have minor effects on the phenotype but may be important to maintaining heritability.

In this context, this work presents a combination of omics data to determine the mechanisms of genotype and phenotype associations involved in rubber tree growth. Using real populations from Hevea breeding programs, association mapping was carried out, and the results were incorporated into RNA-Seq-based coexpression networks and metabolic networks, which were assessed indirectly through the enzymes involved in vegetative growth. By using this established multiomics framework, our study supplies important clues to how the metabolic mechanisms of rubber tree growth are interconnected and suggests novel growth-associated genes for future research on increasing Hevea production.

## 2 Materials and Methods

According to the analysis workflow performed, different molecular layers were investigated in this work (Figure 1). The study started with the identification of the SNPs with the greatest effect on stem diameter (SD) through a GWAS. After selecting these markers, the markers that presented a significant correlation were selected. This entire set of markers was annotated using a transcriptome assembled on the basis of two commercial genotypes that have been widely used in the genetic improvement of the species. Additionally, a weighted gene coexpression network was constructed, from which it is possible to select the functional modules containing the genes identified by the GWAS. An enzymatic network was also built based on the annotation of genes present in the functional modules selected, which supplied insights into the interaction of these enzymes with the studied phenotype.

**Figure 1.**
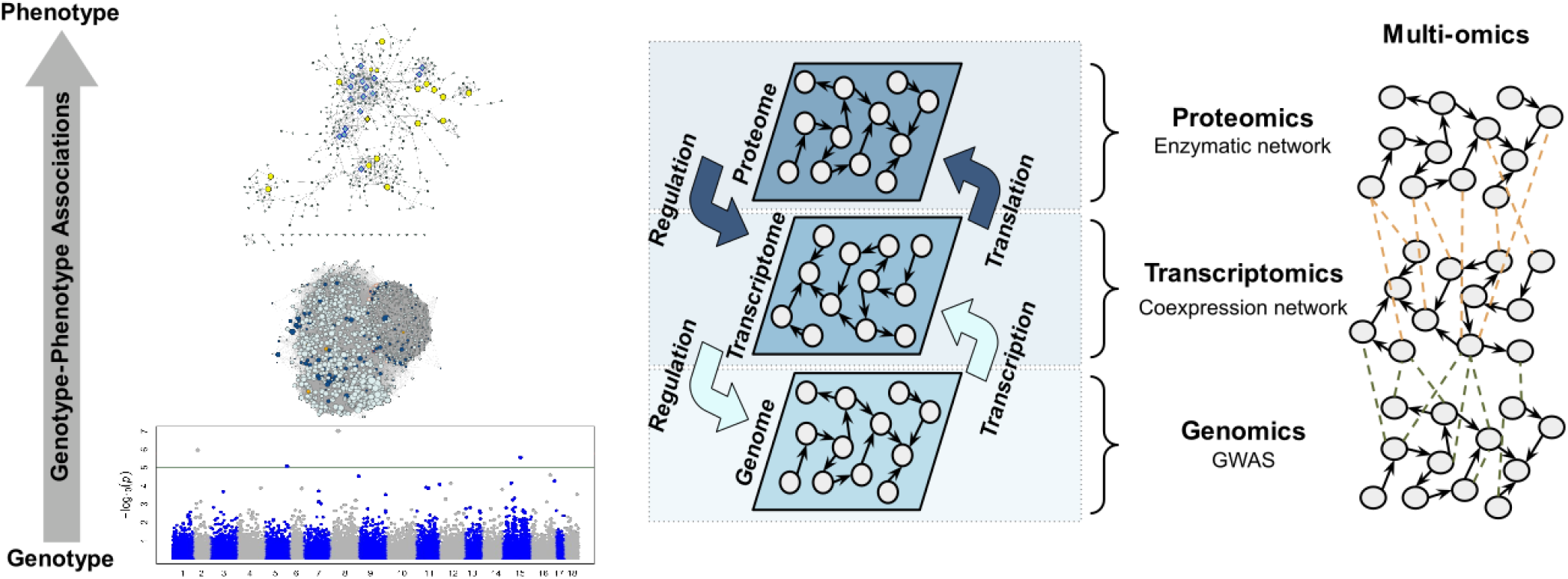
Workflow summarizing the main analyses performed.

### 2.1 Plant Material

For this work, we employed a population composed of 4 test clones (GT1, PB235, RRIM701 and RRIM600) and individuals from crosses between PR255 x PB217 (251 samples), GT1 x RRIM701 (143 samples) and GT1 x PB235 (40 samples) (Souza et al., 2013, 2019; Conson et al., 2018; Rosa et al., 2018). The PR255 genotype was selected because of its early growth and high yield, as well as for being vigorous with stable latex production throughout life (Souza et al., 2013). In contrast, the PB217 genotype presents slow growth but has a rapid increase in latex production in its early years and great potential for long-term performance and yield (Souza et al., 2013; Rosa et al., 2018). These genotypes were planted in random blocks, with four replications of the same genotype grafted on the same plot. This plantation is located in Itiquira, Mato Grosso (MT), Brazil (17° 24′03″ S and 54° 44′53″ W). The GT1 genotype was selected because it is a sterile male and is classified as a primary clone that is tolerant to wind and cold (Shearman et al., 2014). The RRIM701 clone shows vigorous growth and a stem diameter increase after the initial cut (Romain and Thierry, 2011). PB235 has been shown to be a high-yield genotype but is susceptible to panel dryness (Sivakumaran et al., 1988). These two populations (GT1 x RRIM 701 and GT1 x PB 235) were planted in modified blocks that were repeated in four blocks containing two plants of the same genotype per plot with 4 meters of spacing between them. These populations were planted at the Center for Rubber and Agroforestry Systems/Instituto Agronômico (IAC) (20° 25′00″ S and 49° 59′00″ W) in the northwest region of the state of São Paulo (SP), Brazil. All of these genotypes are widely employed in commercial production and used in Brazilian breeding programs, representing the main rubber tree genetic sources in Latin America. Crossing was carried out via open pollination, and paternity was confirmed using microsatellite markers (SSRs).

### 2.2 Phenotypic Analyses

As the main characteristic evaluated in rubber tree genetic breeding (Rao and Kole, 2016), stem diameter (SD) was measured in the selected population during the first 4 years of genotype development. Each plant was individually phenotyped (in centimeters) at a height of 50 cm from the soil in two seasons with contrasting average rainfall (low precipitation and high precipitation), which are considered in Hevea studies as contrasting environments (Chanroj et al., 2017; Souza et al., 2019). The variance caused by the genotypic effects was estimated using the best linear unbiased predictor (BLUP) with the breedR package in R (Munõz and Sanchez, 2017). The linear mixed model was as follows:

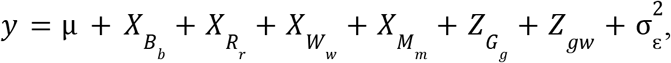

where y is the vector of the phenotypic measures; µis the trait mean; and *X*_*B*_, *X*_*R*_, *X*_*W*_ and *X*_*M*_ are the incidence matrices for the fixed effects of blocks (*b*), replicates (*r*), water levels (*w*) and month of the measurement (*m*), respectively. *Z*_*G*_ and *Z* are the incidence matrices of random effects for genotypic effects (*g*) and genotype x environment interactions (*gw*), respectively, and 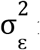 is the residual variance. The significance of random effects was estimated by a likelihood ratio test (LRT) with a significance level of 0.05. We estimated the broad heritability (*H*^2^) for genotypic means using the following equation:

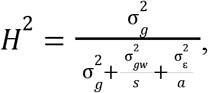

Where 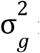 is the genotypic variance, 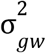 is the variance caused by the environment x genotype interaction, *s* is the number of environments analyzed, 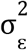 is the residual variance and *a* is number of blocks.

### 2.3 Genotypic Analyses

The extraction of genomic DNA was performed according to Souza et al. (2013) and Conson et al. (2018). Genotyping-by-sequencing (GBS) libraries were prepared from genomic DNA using the method proposed by Elshire et al. (2011). Initially, the genomic DNA of each sample was digested using the methylation-sensitive enzyme EcoT22I to reduce the genomic complexity. The resulting fragments of each sample were linked to specific barcodes and combined in pools. These fragments were amplified by PCR and sequenced. Sequencing of the PR255 x RRIM217 population was performed using the Illumina HiSeq platform, and sequencing of the GT1 x RRIM701 and GT1 x PB235 populations was performed with the GAIIx platform (Illumina Inc., San Diego, CA, United States). Processing of the GBS data from both experiments was carried out at the same time. SNPs were identified with TASSEL GBS 5 software (Glaubitz et al., 2014) using the following parameters: (i) k-mer size of 64 bp; (ii) minimum read quality (Q) score of 20; and (iii) minimum locus depth of 6 reads. Reads were aligned with the rubber tree reference genome proposed by Liu et al. (2020a) using Bowtie2 version 2.1 software (Langmead and Salzberg, 2012) with the very sensitive option. We only kept the biallelic markers selected with the VCFtools program (Danecek et al., 2011). Using snpReady software (Granato and Fritsche-Neto, 2018), SNPs with more than 20% missing data and minimum allele frequency (MAF) <0.05 were filtered out. Imputation was performed using the k-nearest neighbor imputation (kNNI) algorithm (Hastie et al., 2017). LD estimations were calculated with the ldsep R package (Gerard, 2020) based on the squared Pearson correlation (R^2^). For linkage decay investigation, we created a scatter plot of R^2^ against the chromosomal distances, considering an exponential decay (*y* = *a* + *be*^(*cx*)^) (Ranc et al., 2012) created with a nonlinear least squares regression model using R software.

### 2.4 GWAS

GWAS were performed using the Fixed and random model Circulating Probability Unification (FarmCPU) method implemented in the FarmCPU R package (Liu et al., 2016). The kinship matrix and the first two principal components (PC1 and PC2) from a principal component analysis (PCA) were used as covariables in the mixed linear model to control the effects caused by the population structure (Challa and Neelapu, 2018). The significance threshold used for the association mapping was calculated based on 30 SD permutations and a 95% quantile value. Additionally, we expanded the set of putatively associated markers through LD. Considering a minimum R^2^ of 0.7, we created a set of GWAS LD-associated markers (snpsLD), which was used for modeling an LD network with the igraph R package (Csardi and Nepusz, 2006).

### 2.5 Transcriptome

To estimate rubber tree gene expression, RNA-Seq data from RRIM600 and GT1 clones (Mantello et al., 2019) were used. From 6 months of age, these plants were transferred to a growth chamber at a temperature of 28 °C with a 12-hour photoperiod and were irrigated every 2 days for a period of 10 days. After this period, the plants were subjected to cold stress by changing the chamber temperature to 10 °C for 24 hours, with the leaf tissues being sampled at 0 hours (control), 90 min, 12 hours and 24 hours after exposure to the stress. RNA was extracted from the leaves of three biological replicates using the lithium chloride protocol (Dusotoit-Coucaud et al., 2009). From the total RNA, a cDNA library was built using the TruSeq RNA Sample Preparation Kit (Illumina Inc., San Diego, CA, USA). The 24 samples (three replicates per sample at each time) were randomly pooled (4 samples per pool) and grouped using the TruSeq Paired-End Reads Cluster Kit on the cBot platform (Illumina Inc., San Diego, CA, USA). The cDNA libraries were posteriorly sequenced on the Illumina Genome platform Analyzer IIx with a TruSeq kit with 36 cycles (Illumina, San Diego, CA, USA) for 72 bp paired-end reads.

RNA-Seq barcodes were removed from FastQ files using Fastx-Tookit (http://hannonlab.cshl.edu/fastx_toolkit/index.html), and raw reads were filtered using the program NGS QC Toolkit 2.3 (Trivedi et al., 2014), keeping only sequences with a minimum Q-score of 20 across at least 70% of the sequence length. The filtered sequences were combined with bark reads (Mantello et al., 2014) and mapped to the reference genome of *H. brasiliensis* (Tang et al., 2016) using the HISAT2 aligner (Kim et al., 2015). The alignment was ordered and assembled using SAMtools (Li et al., 2009) and Trinity (Grabherr et al., 2011) software, respectively. *H. brasiliensis* scaffolds (Tang et al., 2016) were submitted for ab initio annotation using the Maker-P (Campbell et al., 2014) tool. The Trinity assembled transcripts and the Maker-P annotations were combined with nonredundant *H. brasiliensis* ESTs in the NCBI database (August 2016) and used as a database for aligning assemblies against the *H. brasiliensis* genome (Tang et al., 2016) with the PASA *v2.0* pipeline (Haas et al., 2003) after removing redundant alternate splicing data. The obtained transcripts were filtered with a minimum size of 500 bp and evidence of transcription; we excluded sequences that were only predicted by ab initio genome annotation and with high identity for nonplant transcripts. To estimate the physical position of these sequences across Hevea chromosomes, we performed comparative alignments of these transcripts against the *H. brasiliensis* genome proposed by Liu et al. (2020a) using BLASTn (Johnson et al., 2008). The annotation of these transcripts was performed using the Trinotate v3.2.1 program (Haas, 2015) and SwissProt database (downloaded in February 2021) (Boeckmann et al., 2003).

### 2.6 Gene-Associated Markers

The analysis of candidate genes in QTL regions was performed based on transcript annotations. Candidate genes for the phenotypic variation of GWAS-discovered SNPs were considered by using the first transcripts positioned in the upstream and downstream regions of these markers. In addition to the SNPs significantly associated with the phenotype discovered by the GWAS, which we will call snpsGWAS here, we also searched for candidate genes in the neighboring snpsLD. The GO terms associated with these annotations (snpsGWAS and snpsLD) were investigated using REVIGO (Supek et al., 2011). The genomic regions of the phenotypically associated SNPs discovered in this work were compared with the QTLs discovered by Conson et al. (2018) from the mapping population GT1 x RRIM701. For this analysis, the sequences underlying the QTLs (Conson et al., 2018) were aligned to the reference genome of Liu et al. (2020a) using BLATn. Alignments with identity above 90% and with the largest coverage area were selected (minimum e-value of e-10). Based on the position of this alignment in relation to the reference genome of Liu et al. (2020a), a representation of the 18 chromosomes of *H. brasiliensis* was made using snpsGWAS, snpsLD and QTLs (Conson et al., 2018) using the MapChart program v.2.2 (Voorrips, 2002).

### 2.7 Coexpression Networks

For modeling coexpression networks, we used RNA-Seq count data grouped into transcript clusters through PASA v2.0 software (Haas et al., 2003). Only transcripts with at least 10 counts per million (CPM) were retained and normalized with a quantile-based approach implemented in the edgeR package in R (Robinson et al., 2010). Weighted gene correlation analysis (WGCNA) was performed using the WGCNA R package (Langfelder and Horvath, 2008) together with Pearson correlation coefficients. A soft thresholding power β-value was estimated for fitting the network into a scale-free topology, and a topological overlap measure (TOM) for each gene pair was used for building a dissimilarity matrix and for performing unweighted pair group method with arithmetic mean (UPGMA) hierarchical clustering. The best clustering scheme was defined using a variable height pruning technique implemented in the Dynamic Tree Cut R package (Langfelder, et al., 2008). The groups containing genes associated with snpsGWAS were used to model a specific coexpression network using the igraph R package (Csardi and Nepusz, 2006) with Pearson correlation coefficients (minimum R value of 0.5), where we calculated the hub scores for each gene considering Kleinberg’s hub centrality scores (Kleinberg, 1999).

### 2.8 Metabolic Network Modeling

From the annotations performed for genes surrounding the snpsGWAS and the snpsLD, we retrieved the enzyme commission (EC) numbers and investigated the related metabolic pathways using the Kyoto Encyclopedia of Genes and Genomes (KEGG) database (Kanehisa and Goto, 2000). All the *H. brasiliensis* metabolic pathways with enzymes related to snpsGWAS and snpsLD were retrieved and used to model a metabolic network using BioPython v.1.78 (Cock et al., 2009). From the created network, we evaluate the following topological properties: (i) degree (Barabási and Oltvai, 2004), (ii) betweenness centrality (Brandes, 2001), (iii) stress (Brandes, 2001), (iv) short path length value (Watts and Strogatz, 1998), and (v) neighborhood connectivity (Maslov and Sneppen, 2002), using Cytoscape v3.8.2 (Shannon et al., 2003). The network was also categorized regarding its community structure, with the enzymes organized into modules using the HiDeF algorithm (Zheng et al., 2021).

## 3 Results

### 3.1 Phenotypic and Genotypic Analyses

The SD values were adjusted according to the mixed model from which the BLUPs were extracted for further analysis (Supplementary Table 1). All fixed and random effects showed significant effects under the LRT test (p <0.01). The estimated variances were 4.56, 0.0001 and 26.69 for the genotype (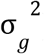), genotype x environment interaction 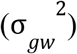 and residual 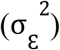 effects, respectively. The experimental design was confirmed to show normality of the residual variance based on the quantile-quantile graph (Q-Q plot) (Supplementary Figure 1). The estimated heritability (*H*^2^) in the entire population was 0.55, which is close to those values found in previous studies on the species (Gonçalves et al., 1999; Chanroj et al., 2017).

The identification of SNPs was carried out using all 437 individuals. By employing the TASSEL pipeline, we produced 363,641 tags, which were aligned with the Hevea reference genome, producing an alignment rate of ∼84.78%. We identified a total of 107,466 SNPs, which were filtered, resulting in a total of 30,266 high-quality markers (∼28.16%), with an imputation rate of ∼6.74%.

This filtered SNP dataset was used for PCA, with 18.33% and 2.61% of the variance explained by the first two main components, respectively (Supplementary Figure 2). Although high LD decay was observed (Supplementary Figure 3A), we also assessed the LD decay rate only in the regions containing transposable elements (TEs) (Supplementary Figure 3B), which was higher.

### 3.2 RNA-Seq Analyses

A total of ∼530 million and ∼633 million paired-end (PE) reads were obtained for the RRIM600 and GT1 genotypes, respectively. After quality filtering, we obtained ∼933 million PE reads for assembling the transcripts through Trinity software. We identified 104,738 transcripts ranging from 500 bp to 22,333 bp (average transcript size of 1,874 bp and N50 of 2,369 bp) that were related to 49,304 genes . In total, 82,629 transcripts (78.89%) could be annotated using the Swiss-Prot database. We were able to associate Gene Ontology (GO) categories with 81,095 transcripts (77.42%) and metabolic pathways from the KEGG database with 74,668 transcripts (71.29%). A total of 11,150 different proteins could be associated with the estimated set of genes for rubber trees, with a high incidence of TEs; the retrovirus-related Pol polyprotein from transposon RE1 (RE1) (4.45%) and the retrovirus-related Pol polyprotein from transposon TNT 1-94 (TNT 1-94) (2.80%) were the most pronounced categories.

### 3.3 Genome Wide Association Study

With the FarmCPU method and the selected covariates, we were able to observe satisfactory adherence to the association mapping results (Figure 2A). Four snpsGWAS were identified on chromosomes 2, 5, 8 and 15 (Figure 2B). The MAFs of the snpsGWAS ranged from 10 to 45%, with additive heritabilities ranging from 2 to 9% and additive effects ranging from -1 to 0.84 cm (Table 1). To assess all markers associated with SD, we expanded the set of significantly associated markers by means of LD tests on the total set of SNPs. A total of 181 snpsLD were found and showed a correlation greater than 0.7 with the snpsGWAS (Supplementary Figure 4). snpsLD are distributed on the 18 chromosomes of the rubber tree (Figure 3), flanking previously described QTLs (Conson et al., 2018). We were able to identify SNPs with distances of approximately 40 bp in the QTL regions (Figure 3).

**Figure 2.**
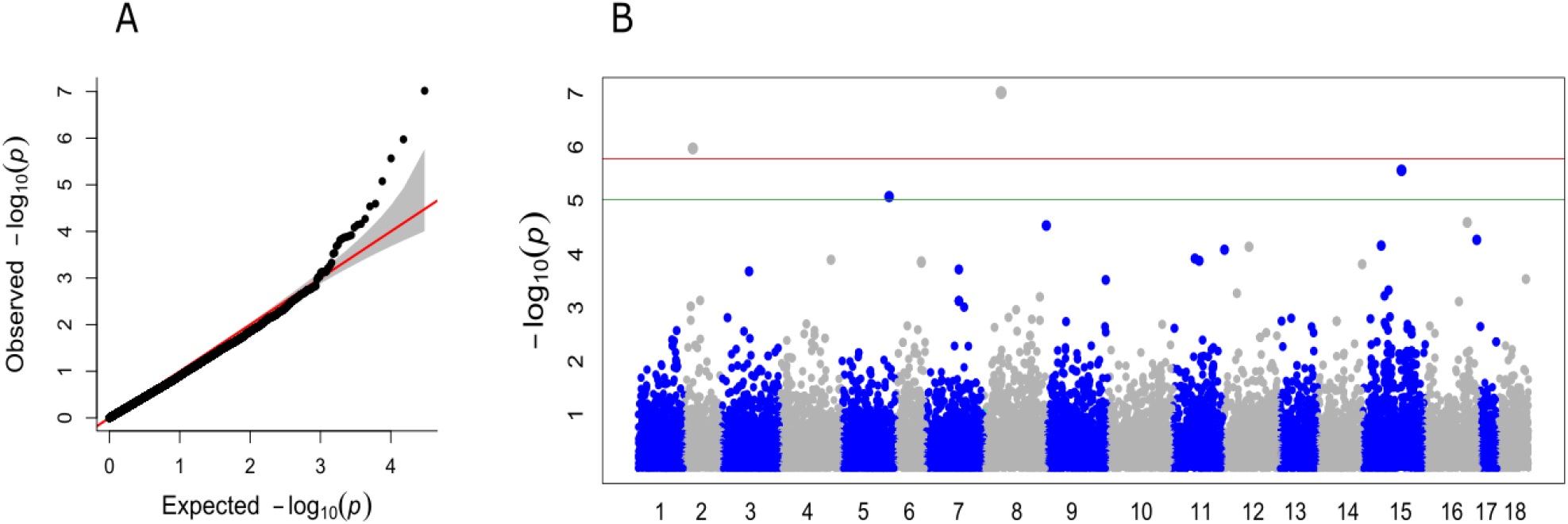
(**A**) Quantile-quantile plot for the broad genomic association model (GWAS), with the inclusion of the first main component (PC1) as a covariate. (**B**) Manhattan plot for the GWAS. The X axis shows the chromosomes containing the discovered markers in their respective positions. The Y axis shows the log (p-value) of the association. The red line represents the threshold obtained based on the data, and the green line represents the Bonferroni-corrected threshold of 0.05.

**Figure 3.**
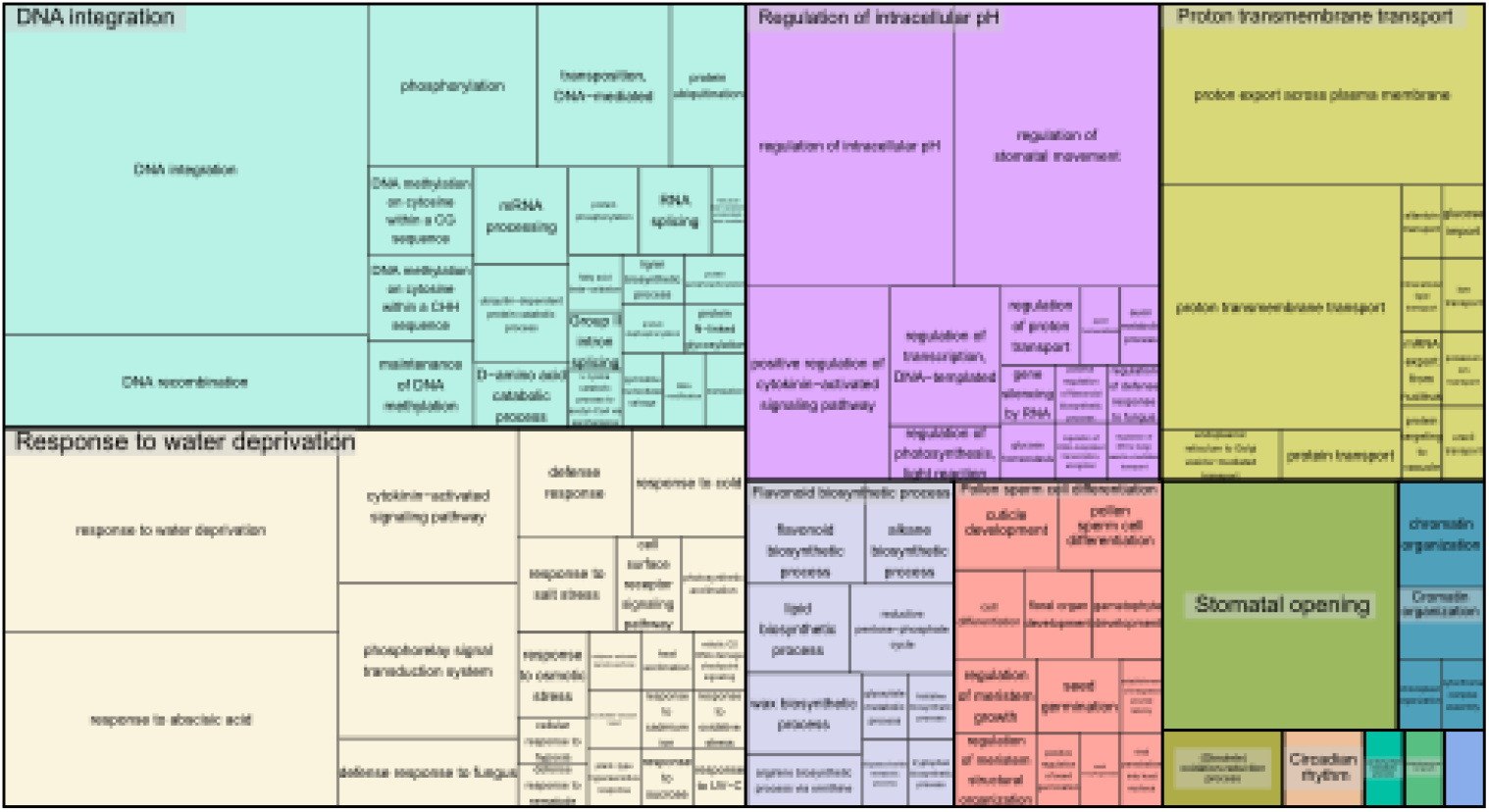
Treemap representing the biological processes for the GO terms of the annotated SNPs.

**Figure 4.**
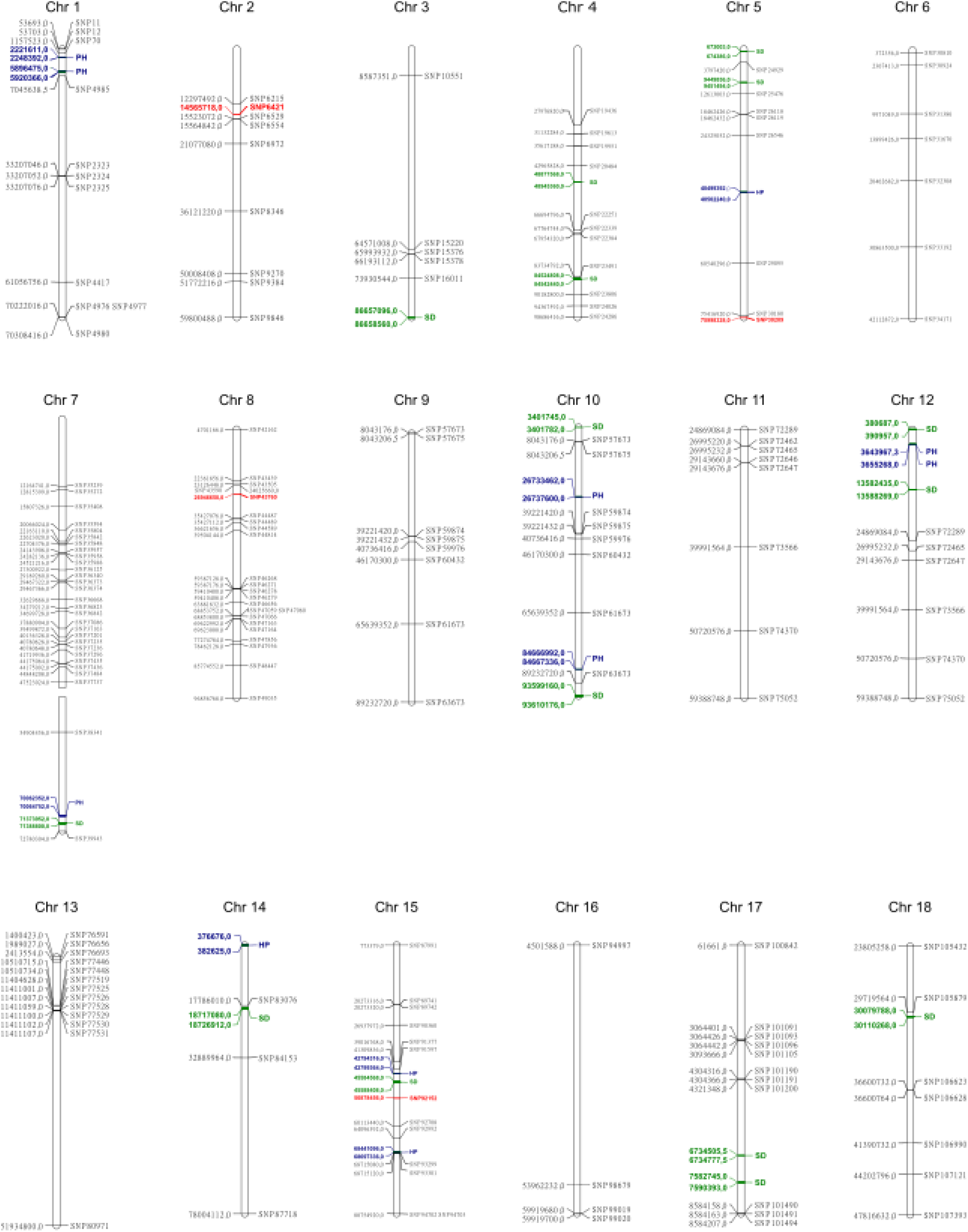
Physical position of snpsGWAS in red, snpsLD in black and QTLs discovered by Conson et al. (2018). The QTLs for plant height (PH) are in blue and those for stem diameter (SD) are in green.

**Table 1.**
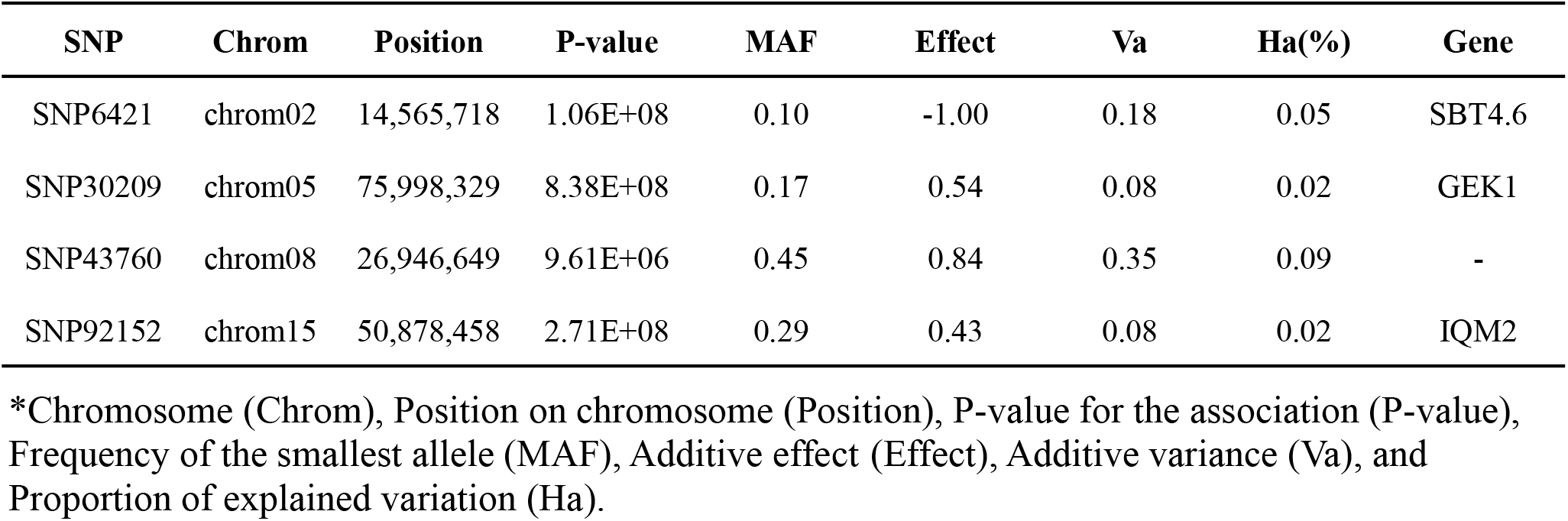
SNPs identified through the GWAS model.

To infer the associations between the set of SNPs (snpsGWAS and snpsLD) and expressed genomic regions, we performed comparative alignments of the transcripts assembled to the rubber tree chromosomes. SNPs were assigned to the first genes that were downstream and upstream of their location with an average distance of 7 kbp (Supplementary Table 2). Among the snpsLD, genes related to the transcription of important proteins involved in different stresses were found, such as TNT 1-94, receptor-like protein EIX2, integrin-linked protein kinase 1, U1 small nuclear ribonucleoprotein 70 kDa, histidine-containing phosphotransfer protein 2, rhomboid-like protein 14, and mitochondrial and threonine-protein kinase STN7. The annotation of the set of SNPs putatively associated with SD showed major biological processes related to DNA integration, response to water deprivation, regulation of intracellular pH, proton transmembrane transport, stomatal opening, flavonoid biosynthetic process, pollen sperm cell differentiation, oxidation-reduction process, circadian rhythm, carbohydrate metabolic process, multidimensional cell growth and chromatin organization (Figure 3).

### 3.4 Gene Coexpression Network

Of the 104,738 transcripts, 30,407 were selected for modeling a gene coexpression network using the WGCNA methodology (Zhang and Horvath, 2005). In such a network, pairwise gene interactions are modeled through a similarity measure, such as the Pearson correlation coefficient employed here. For fitting the network into a scale-free topology, we selected a β power of 9 (scale-free topology model fit with *R*² > 0. 85 and mean connectivity of ∼183.47) and calculated the corresponding dissimilarity matrix through the WGCNA R package. With the network modeled, we combined UPGMA clustering with a variable height pruning technique, enabling the identification of 174 groups, with sizes ranging from 52 to 3,823 genes. The 5 groups containing the genes potentially related to the snpsGWAS were selected (Supplementary Table 3), and a new coexpression network was built including the genes associated with the snpsLD (Figure 5). All these genes formed a unique interaction network with weaker interactions connecting the found groups, which putatively represents the direct and indirect molecular associations with the SD phenotype. For the analysis of all reactions triggered by the genomic regions associated with GWAS, we evaluated this set of 1,528 genes for related GO terms (Figure 6). From the biological process category, we found new GO terms not associated with the genes related to snpsGWAS and snpsLD. These GO terms included defense response, positive regulation of transcription, cell wall organization, photosynthesis, cell division, mitotic cell cycle phase transition, carbon fixation, cell population proliferation, asymmetric cell division, and stomatal closure.

**Figure 5.**
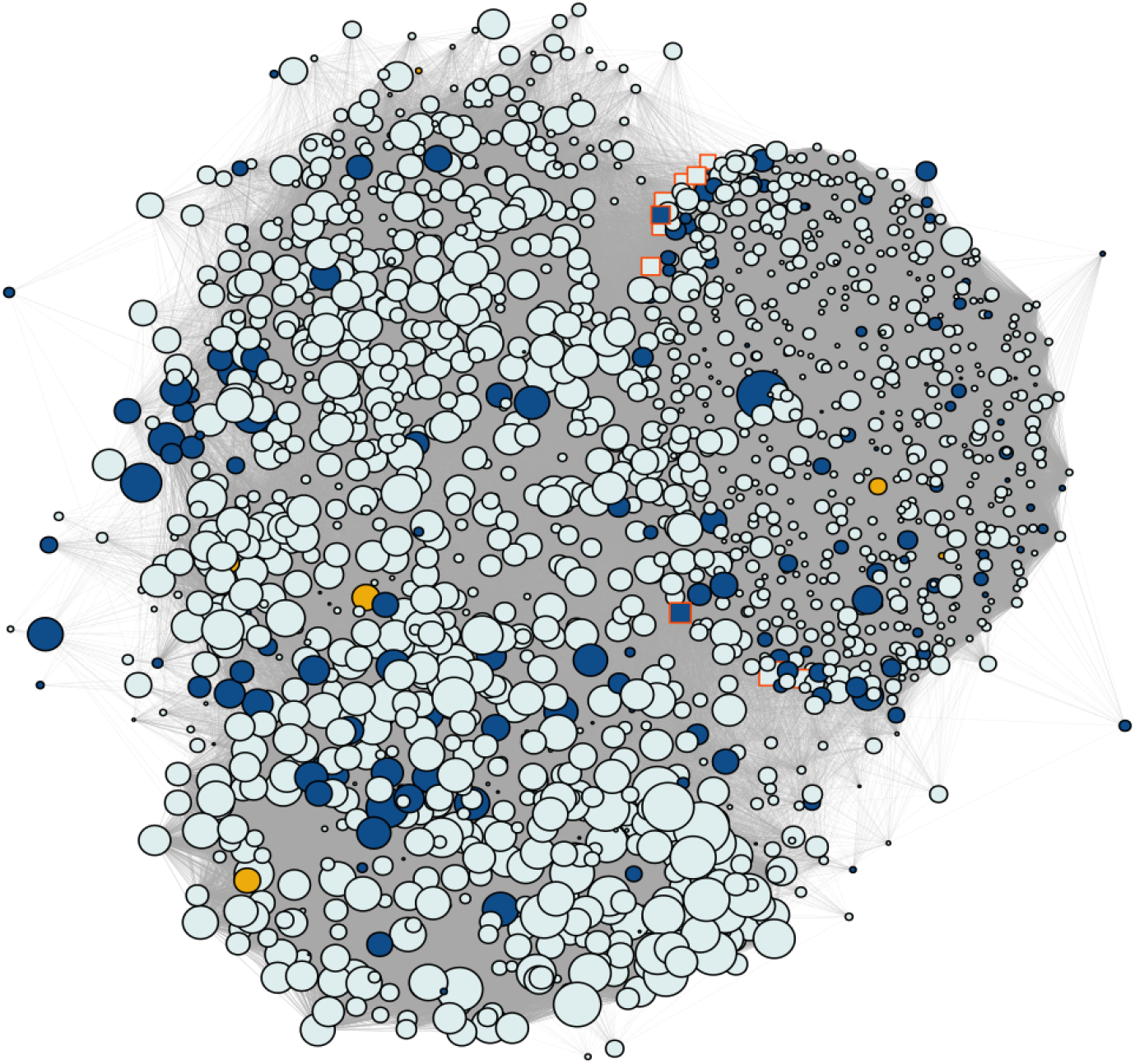
Coexpression network containing the SNP gene modules discovered by GWAS. Yellow shows the genes annotated for the snpsGWAS, blue shows the genes annotated for the snpsLD and gray shows the genes identified in the modules. The highlighted genes with a red border represent the 10 hubs with the most connectivity, while the size of the nodes shows the number of connected genes.

**Figure 6.**
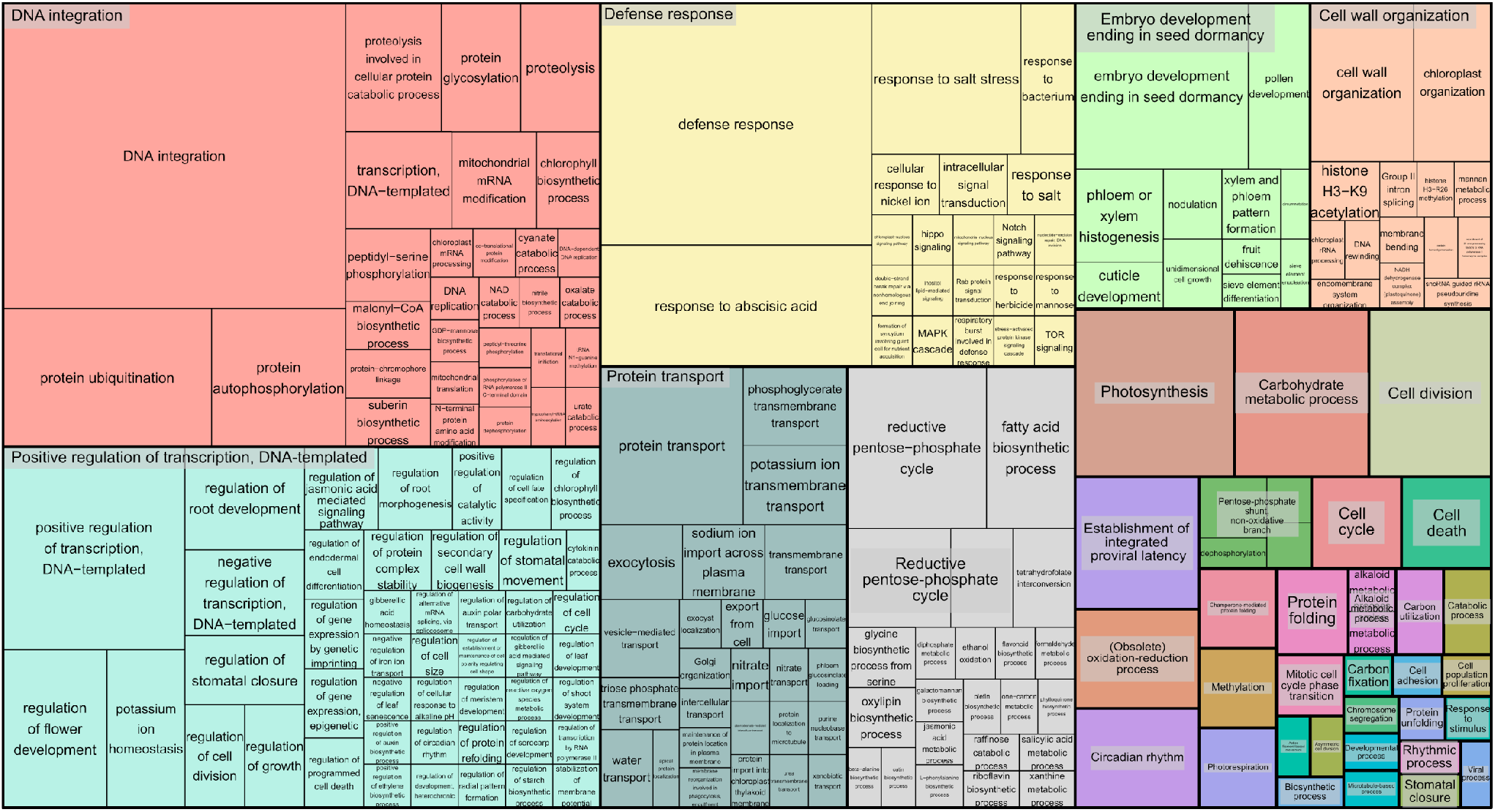
Treemap representing the biological processes for the GO terms of the annotated functional modules.

Regarding the genes found in these modules, as also observed in the general transcriptome profile, we observed a predominance of genes related to the protein retrovirus-related Pol polyprotein from transposon 17.6 (TE 17.6) (2.36%) and TNT 1-94 (1.23%). We also found several genes related to proteins involved in (Supplementary Table 3): (i) plant growth (e.g., MEI2-like 4 and threonine-protein kinase GSO1); (ii) the response to biotic and abiotic stress (e.g., abscisic acid-insensitive 5-like protein 6, transcription factor ICE1, abscisic acid receptor PYL4, transcription factor jungbrunnen 1, transcription factor MYB44, and galactinol synthase 2); (iii) root growth (e.g., alkaline/neutral invertase CINV2, threonine protein kinase IREH1, phospholipase D zeta 1, protein arabidillo 1, regulatory-associated protein of TOR 1, agamous-like MADS-box protein AGL12, and omega-hydroxypalmitate O-feruloyl transferase); (iv) the hormone abscisic acid (ABA) pathway; and (v) the light acclimatization process (e.g., GATA transcription factor 7 and malate dehydrogenase [NADP]). However, the great majority of these identified genes were not overexpressed, with a few exceptions (Supplementary Figure 5). To assess the most influential nodes within the network structure, we evaluated the hub scores of each gene within the network. The first hub gene in this network (PASSA_cluster_140395) was among the snpsLD genes, and the 10 first hubs had many known annotations. The first three hubs that had a known annotation were PASA_cluster_160224, PASA_cluster_87395, and PASA_cluster_140392, showing associations with TEs (Supplementary Table 3).

### 3.5 Metabolic Networks

Due to the clear absence of functional annotations, all the genes identified in the coexpressed modules with a known enzymatic activity relatedness were used for modeling a metabolic network using the KEGG database. In this structure, each enzyme corresponds to a node, and their connections are based on metabolic interactions. Nineteen genes were related to 19 different enzymes present in 28 metabolic pathways (Supplementary Table 4). All these reactions were joined into a unique network structure containing 405 nodes (enzymes) and 1,311 edges (average number of 5.338 neighbors and diameter of 22 nodes) (Figure 7A; Supplementary Figure 6), representing a diverse cascade of mechanisms with putative associations with plant growth. Network topology measurements were performed to identify the most important enzymes in the modeled mechanisms.

**Figure 7.**
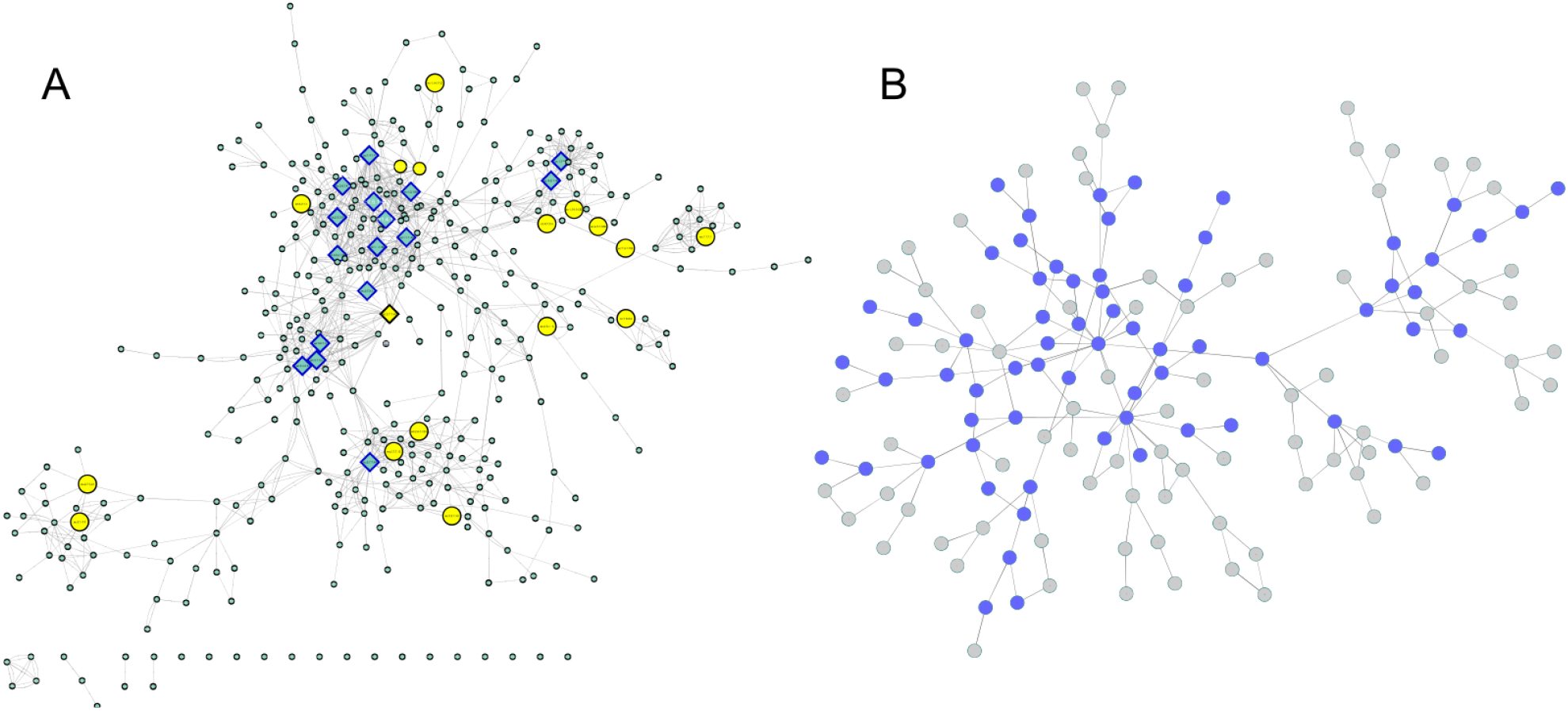
(**A**) Enzyme network. The yellow nodes represent the enzymes discovered in the coexpression modules, and the rectangular nodes indicate the enzymes with the highest centrality values. (**B**) Communities. The blue nodes are represented by communities containing enzymes discovered in the coexpression modules.

From the degree measures for each node (considering in and out connections), we identified 17 outliers (Figure 7A; Supplementary Figure 6), which were considered network hubs. We found enzymes with diverse roles (Supplementary Table 5), such as UDP-sugar pyrophosphorylase (ec: 2.7.7.64) (34 connections), ureidoglycolate amidohydrolase (ec:3.5.1.116) (26 connections) and alanine-glyoxylate transaminase (ec:2.6.1.44) (25 connections). Interestingly, these enzymes were also the ones with the highest values of outdegree, stress and betweenness. Considering only the indegree connections, the top 4 enzymes (also identified among the network hubs) were UDP-sugar pyrophosphorylase (ec: 2.7.7.64) (34 connections), glutamate dehydrogenase (NAD (P) +) (ec: 1.4.1.3) (19 connections), glutamate dehydrogenase (NADP+) (ec: 1.4.1.4) (18 connections) and malate dehydrogenase (oxaloacetate-decarboxylating) (NADP+) (ec: 1.1.1.40) (15 connections). Among the 17 hubs, pyruvate kinase (ec: 2.7.1.40) also presented high values for other centrality measures (betweenness and stress). Additionally, the enzyme threonine synthase (ec: 4.2.3.1) showed the highest short path length value (14.30) and the highest eccentricity value (22), and the glucuronokinase enzyme (ec: 2.7.1.43) showed the highest value for neighborhood connectivity (24).

In addition to these evaluations, the modeled network was also categorized into condensed modules regarding the community structure and enzyme organization (Figure 7B; Supplementary Figure 7; Supplementary Table 6). Using the HiDeF (Zheng et al., 2021) algorithm, 149 communities were identified, containing 4 to 389 enzymes. The community with the highest eccentricity (7) was c1337, which also contained the highest number of enzymes. The community with the highest stress value (255) and betweenness (0.37) was c13340 (with 68 enzymes), and it was among the top 3 communities with the highest eccentricity value (5) (Supplementary Table 7).

## 4 Discussion

The genetic improvement of rubber trees requires a long period of time, with more than 30 years estimated for developing an improved genotype (Gonçalves and Fontes, 2012). Despite the specialized labor required for Hevea phenotyping, its plantation is only possible in vast areas, making the selection process laborious and financially expensive. In this context, the use of MAS can drastically reduce the time and the cost of genetic improvement, especially if implemented in the first years after obtaining the seeds by selecting the target characteristics indirectly through phenotypically associated markers (Xu and Crouch, 2008). As a way of assisting such initiatives, in this work, we identified SNPs associated with SD, and this set of markers can be used as high-priority candidates for MAS, with a high potential of providing greater precision and requiring less time in the selection of superior genotypes.

The main abiotic limitations for the productivity of cultivated plants are excessive salinity, adverse temperatures, and water deficit (Zhu, 2016), and for rubber tree production, water stress and cold are widely described as the most impactful limitations (Ding et al., 2020). Although several studies have investigated the molecular mechanisms of Hevea in cold resistance for its improvement (Cheng et al., 2018; Deng et al., 2018; Mantello et al., 2018), one of the main characteristics evaluated in Hevea breeding programs is SD (Priyadarshan, 2003) due to its versatility in assessing rubber tree productive efficiency (Dijkman, 1951; Goncalves et al., 1984; Chanroj et al., 2017; Conson et al., 2018; Khan et al., 2018; Chen et al., 2020). The use of SD measures can provide insights into phenotypes that can only be measured in specific climate conditions, such as drought resistance (Ohashi et al., 2006; Zhang et al., 2019a), which impacts rubber tree growth (Chandrashekar et al., 1998). Additionally, traits that can only be measured after a certain age of the plant, such as the production of latex and vigor (Dijkman, 1951; Goncalves et al., 1984), can be estimated by SD.

As SD is a quantitative characteristic, the study of the genetic architecture related to this trait is quite complex, considering the high amount of genes and metabolic pathways involved in its definition (Pootakham et al., 2020). Furthermore, the genome of rubber trees encompasses a large number of repetitive regions, reaching approximately 71% of rubber tree genomic content (Tang et al., 2016). The first Hevea reference genome at the chromosome level was only recently published in 2020 (Liu et al., 2020a), and most genomic approaches in the species have been based on highly fragmented sequences and biocomputational estimations (Pootakham et al., 2015; Chanroj et al., 2017; Conson et al., 2018; de Souza et al., 2018; Souza et al., 2019). Only with the advent of molecular biology techniques for reducing genomic complexities during sequencing procedures, such as GBS (Elshire et al., 2011; Poland and Rife, 2012), has it been feasible to generate thousands of SNP markers with high frequency in complex plant genomes (Pootakham et al., 2015). By using a GBS approach combined with a rubber tree chromosome-level reference genome, we characterized a large number of high-quality markers regarding their genomic distribution and LD relatedness, which enabled us to compare our findings with the locations of several QTLs for this characteristic, embracing novel possible causal genes explaining this phenotypic variation.

The large number of SNP markers discovered in this work allowed us to assess the LD throughout the genome of the entire population in a very representative way. As in other studies using arboreal and allogamous species (Peláez et al., 2020), our results showed high LD, which is consistent with previous *H. brasiliensis* results (Chanroj et al., 2017; De Souza et al., 2018). Interestingly, such elevated decay is not constant, and regions with a high density of TEs present a lower level of LD compared to the overall genomic LD. TEs are known as mobile elements due to their ability to change positions along the genome and produce copies of themselves (Singh et al., 2019), mainly in genomic regions with low LD (Stuart et al., 2016; Choudhury et al., 2019), as was observed in this work (Supplementary Figure 3). As stated by Choudhury et al. (2019), we also believe that there are two main reasons for this observation: (i) TEs alter the genetic architecture of the chromosome by decreasing the recombination rate in its vicinity; and (ii) TEs accumulate in these regions due to the low recombination rate that occurs in these locations.

Several studies have been developed to characterize SD QTLs (Souza et al., 2013; Conson et al., 2018; Rosa et al., 2018); however, these studies are limited to the biparental populations employed (Myles et al., 2009). With the use of genetically diverse populations, GWAS approaches use the historical links between different genotypes, capturing more genetic diversity through a broader set of markers that would be neglected in association maps (Kulwal, 2018). When we are unable to identify the expected segregation ratios in markers from biparental progenies, these regions, even those close to important QTLs, are often discarded along with their associated QTLs (Kulwal, 2018). In this context, GWAS approaches have been suggested as a powerful tool for overcoming such limitations, which are intensified in species such as *H. brasiliensis*, in which there are great difficulties in obtaining mapping populations.

### 4.1 GWAS

To date, only one study employing GWAS has been described in the literature for *H. brasiliensis*. Using a population of 170 individuals genotyped with 14,155 SNP markers by capture probes (Shearman et al., 2014), Chanroj et al. (2017) tested four association models. The authors could associate two SNP markers with latex production (one for the rainy season and the other one for the drought season) and two others with SD (also separated by rainy and drought seasons). According to Conson et al. (2018), the rubber tree populations planted in the escape areas are under water stress at all times, despite the differences in water regime across seasons. Due to such observations and the Brazilian climate, we performed our analyses without making this distinction. In this way, we identified 4 SNPs associated with SD, which were annotated following an RNA-Seq-based approach.

Different from establishing a genomic window surrounding these markers and performing comparative alignments against plant databases (Chanroj et al., 2017; García-Fernández et al., 2021), we used an assembled transcriptome for the association of the snpsGWAS. This step was performed mainly because of the absence of available data for several neglected species in public databases (Schaefer et al., 2018), such as *H. brasiliensis*. Moreover, transcriptome assemblies are a way of categorizing a broader range of important genes found under stress conditions (Valdés et al., 2013; Wei et al., 2021), as already reported by other Hevea studies (Ahn et al., 2017; Mantello et al., 2019). Additionally, because of the recent availability of the Hevea genome (Liu et al., 2020a), more studies are required for complete and accurate gene categorization. By coupling the transcriptome assembly with GWAS, we could associate the 3 candidate genes identified by the snpsGWAS, which were annotated and had their expression profile estimated in two different genotypes, including in the population used here, and in specific stages of the plant development and physiology. As pointed out by Schaefer et al. (2018), this type of strategy provides associations not only with growth but also with resistance to abiotic stress. The genes identified flanking the snpsGWAS were interpreted according to their biological function and their metabolic context (Watanabe et al., 2017), suggesting their potential relationships in defining the phenotype.

The SBT4.6 gene (Table 1), identified based on the snpsGWAS, belongs to the subtilisin-like protease family, whose members are involved in general protein turnover and regulatory processes and in mechanisms of resistance to biotic and abiotic stresses (Tian et al., 2005; Budič et al., 2013; Figueiredo et al., 2018). Under normal conditions, mutants for this gene do not show obvious changes in the normal growth of the plant, so there is still a need for further investigations regarding this gene in the development of the plant (Rautengarten et al., 2005). Although this gene is not clearly involved in plant growth under normal conditions, we suggest that it may be indirectly related to this characteristic. Two other genes associated with snpsGWAS show evidence of a relationship with abiotic stresses. In addition to showing an increasing additive effect of 0.54 cm in the SD for a specific genotypic class (Table 1), SNP30209 was in the vicinity of a genomic region containing a candidate gene for GK1. In experiments carried out with *Arabidopsis thaliana*, GK1 showed a behavior of D-aminoacyl-tRNA deacylase, which is important for protecting the plant against the toxicity of D-amino acids (Wydau et al., 2007), which, when present in the soil, can have effects on plant growth in different ecosystems, whether managed or not. These compounds can act in different ways on root and stem growth, with D-serine, D-alanine and D-tyrosine being the strongest growth inhibitors, while others, such as D-lysine, D-isoleucine, D-valine, D-asparagine and D-glutamine, act as milder inhibitors (Vranova et al., 2012). Another associated gene was IQM2, which contains a domain for the IQM2 protein. Such a protein belongs to a calmodulin-binding family protein and has strict involvement in the response to biotic and abiotic stress (Wan et al., 2012).

Despite the unquestionable importance of GWAS methods, the practical application of these findings in MAS for the selection of several complex characteristics is limited due to the low heritability associated with these markers (Bogardos, 2009). Considering this fact, we also investigated associated genomic regions, which may be jointly involved in phenotype definition (Yuan et al., 2012). Several statistical methods are used to identify genomic associations, such as multifactor dimensionality reduction (Ritchie et al., 2001), LD (Wu et al., 2008) and entropy-based statistics

(Dong et al., 2008). In this work, we employed SNP correlations, which led us to already establish QTL positions (Conson et al., 2018), showing the robustness of this method. These newly identified markers may reveal genes that would be overlooked by conventional GWAS approaches. In addition to MAS, other important tools for the genetic improvement of various plant species have been developed, such as iRNA (Zang et al., 2017) and CRISPR (Jaganathan et al., 2018), which have shown enormous potential for breeding strategies in recent years (Kalunke et al., 2020; Liu et al., 2020b). However, these approaches require the definition of target genes and their interactions, which might be estimated through coexpression and metabolic networks. In this way, to provide a deeper investigation into the metabolic activities of the genes associated with the snpsGWAS and snpsLD, we modeled complex networks to investigate their interactions and provide insights into the definition of the SD quantitative trait (Kosová et al., 2015; Tam et al., 2019), decreasing the variability of the indirectly selected phenotype and accessing other omics layers. The multiomics approaches employed here can contribute to a better understanding of the molecular mechanisms that are important to the vegetative growth of rubber trees, opening new perspectives for deeper genomic studies.

### 4.2 Multiomics

Quantitative traits are strongly affected by environment x genotype interactions (Nguyen et al., 2019). Genotypes with a greater capacity to resist these abiotic factors have a greater capacity to grow and develop under these stresses (Mantello et al., 2019). Understanding all the molecular biological levels that confer such a resistance to these specific genotypes requires the integration of multiple omics approaches, such as genomics, transcriptomics, proteomics and metabolomics. Multiomics approaches have as their main objective the integration of data analysis of different biological levels for a better understanding of their relationships and the functioning of a biological system as a whole (Joyce and Palsson, 2006). The use of joint approaches benefits from including all relevant parts that integrate the analyzed biological system (Zhang et al., 2010). Studies that integrate the discovery of QTLs with other omics have used genetically well-studied agricultural crops such as corn (Jiang et al., 2019) and, more recently, tree species such as citrus (Mou et al., 2021).

To provide deeper insights into the molecular basis of the evaluated phenotype, we extended the selected set of SD-associated SNPs with data from transcriptomics using complex network methodologies. These methodologies have revolutionized research in molecular biology because of their capability to simulate complex biological systems (D’haeseleer et al., 2000; Liu et al., 2020c) and infer novel biological associations, such as regulatory relationships, metabolic pathway inferences and annotation transference (Rao and Dixon, 2019). In *H. brasiliensis*, coexpression network methods have already revealed genes involved in different environmental or stress conditions and are a powerful tool for profiling rubber tree samples (Hurtado Páez et al., 2015; Sathik et al., 2018; Mantello et al., 2019; Deng et al., 2020; Ding et al., 2020). Such studies in rubber trees are still incipient and have not yet been coupled with breeding strategies for the genetic improvement of the species. Starting from RNA-Seq-based data, we could associate our GWAS results with expression profiles from important *Hevea* genotypes, incorporating our results into a complete set of molecular interactions estimated through the WGCNA approach. Using this strategy, we can infer biological functions for genes present in the same network module, as these genes probably exert correlated functions (Child et al., 2011). This is the first initiative that proposes the integration of GWAS and coexpression networks in rubber trees to identify genes with great potential to be used in MAS.

The transcriptome used for annotation and construction of the coexpression network showed a large number of TEs, which are indeed present in large amounts in plant genomes (Matsunaga et al., 2015). In addition, these TEs were also found to be abundant in the selected functional modules, with TE 17.6 and TNT 1-94 being the most prominent. These TEs have already been described as being involved in gene expression, responses to external stimuli and plant development (Kashkush et al., 2003; Matsunaga et al., 2015; Traylor-Knowles et al., 2017; Tran and Choi, 2020). In rubber trees, TEs may be related to the differential expression observed in some commercial clones, affecting important processes such as rubber production (Wu et al., 2020). As pointed out by Wang et al. (2020), the identification of TEs associated with functional genes related to important characteristics suggest that they can be used as molecular markers in MAS, contributing significantly to the genetic improvement of woody trees. In this sense, our findings supply a wide range of genomic resources for breeding. In the selected coexpression network, the most abundant elements were also TE 17.6 and TNT 1-94.

In the coexpression module with the largest number of genes, we were able to identify many genes related to plant growth, such as the protein MEI2-like 4 (ML4), which is a substrate for putative TOR, the main regulator of cell growth in eukaryotes (Anderson and Hanson, 2005), representing an extremely important molecule in meiotic signaling (Watanabe et al., 1988). In this module, we also identified the proteins alkaline/neutral invertase CINV2 (CINV2) and LRR receptor-like serine/threonine-protein kinase GSO1 (GSO1), which are related to root growth and endoderm. The invertase enzyme (INV) is one of only two enzymes capable of catabolizing physiological carbon, together with the sucrose synthase enzyme (SUS); thus, most of the plant biomass is indispensable for normal growth, and the loss of these genes slows plant growth (Barratt et al., 2009). According to Racolta et al. (2014), the GSO1 protein works together with GSO2 for the intracellular signaling of the plant, positively regulating cell proliferation, the differentiation of root cells and the identity of stem cells.

In the other functional modules, we identified several proteins involved in abiotic stress, such as transcription factor ICE1 (SCRM), an upstream transcription factor that regulates cold CBF gene transcription, improving plant tolerance to freezing (Chinnusamy et al., 2003). The regulatory-associated protein of TOR 1 (RAPTOR1) presents itself as a TOR regulator in response to osmotic stress (Mahfouz et al., 2006). The transcription factor jumgbrunnen 1 (JUB1), which delays senescence, also confers resistance to abiotic stress, such as heat shock, and resistance to high levels of intracellular H_2_O_2_ (Wu et al., 2012). Protein galactinol synthase 2 (GOLS2) plays an important role in the response against drought and cold stresses (Taji et al., 2002). The protein E3 ubiquitin-protein ligase PUB23 (PUB23), which responds quickly to water stress (Cho et al., 2008) and biotic stress, and the protein glucan endo-1,3-beta-glucosidase (HGN1) have been reported in *H. brasiliensis* and participate in a defense response against fungi (Galicia et al., 2015). In addition to these proteins produced in response to a given stress, genes involved in the maintenance and development of vegetative parts important for the development of the plant under a given stressful condition, such as constant drought, were identified, including arabidillo 1 protein (FBX5), which is related to the development of the roots (Coates et al., 2006), and Agamous-like MADS-box protein AGL12 (AGL12) (Tapia-López et al., 2008). These results confirm the involvement of genes identified by GWAS and other genes identified in functional modules in the investigated characteristic definition. We can also relate the region of the SNP43760 marker, which has no known annotation, to QTLs involved in resistance to environmental factors, since the functional module containing these genes is related to this process.

Finally, a metabolic network for the enzymes found in this data set was constructed to identify the main metabolic pathways involved in the growth process of the rubber tree. The metabolites produced in cells can be understood as a bridge between the genotype and the phenotype. A clearer understanding of the relationship between these enzymes, such as by identifying the main enzymes present in the network, is essential for maintaining the properties of this network and thus preserving these relationships. The network built in this work shows some disconnected enzymes because the reactions that connect them with the other enzymes in the network have not yet been elucidated.

We identified UDP-sugar pyrophosphorylase (USP) as the hub of this enzyme network; this enzyme indicated to be an enzyme of great importance in the network, as it presented the highest degree value (Barabási and Oltvai, 2004). It also showed the highest out-degree value, which represents the number of connections directed from this node to the other nodes in the network. This enzyme is very conserved in plants (Geserick and Tenhaken, 2013). Evidence indicates a high affinity of USP for acid-1-phosphate (UDP-GlcA-1-P), a substrate of the myo-inositol oxygenase (MIOX) pathway for UDP-GlcA (Geserick and Tenhaken, 2013). USP can also convert different types of sugar-1-phosphatates into the UDP sugars that make up polymers and glycerols in plant cell walls (Geserick and Tenhaken, 2013). USP is found in a single copy in Arabidopsis, and mutants for this gene are lethal (Geserick and Tenhaken, 2013), as the pollen that carries this mutation does not develop normally (Schnurr et al., 2006; Geserick and Tenhaken, 2013). Knock-down mutants also show impaired vegetative growth due to deficiency in sugar recycling (Geserick and Tenhaken, 2013). The enzyme glutamate dehydrogenase (NAD (P) +) (GDH) appeared in the enzymatic network containing a high degree of indegree. GDH catalyzes the deamination of glutamate using NAD as a coenzyme and releases 2-oxoglutarate and ammonia when there is little carbon (Fontaine et al., 2006). Participating in the response to various stresses, including drought and the presence of pathogens, their expression levels are regulated according to the intensity of the stress (Restivo, 2004), increasing the capacity of resistance to stress and the acquisition of biomass by the plant (Qiu et al., 2009; Tercé-Laforgue et al., 2015). The pyruvate kinase enzyme was shown to be central in the integration of its components, presenting a higher value of betweenness centrality (Brandes, 2001), indicating an important control function of this enzyme in the network, since this measure indicates elements in the network that join communities. In addition to this enzyme being important for the integration of the components in the metabolic network, this enzyme also presents itself as important in the dissemination of information among the elements present in the metabolic network, since it presented a higher stress value (Brandes, 2001), which indicates the shortest path between two random nodes in the network. This enzyme is a key element in the regulation and adjustment of the glucose metabolic pathway (Ambasht and Kayastha, 2002; Cai et al., 2018). Pyruvate kinase catalyzes the irreversible transfer of the high-energy phosphate group from phosphoenopyruvate to ADP, synthesizing ATP (Ambasht and Kayastha, 2002). Another important enzyme for the dissemination of information within the network was threonine synthase (thrC), which showed a higher value for the short path length (Watts and Strogatz, 1998) and eccentricity, which indicates the maximum number of nodes necessary for the information to reach all nodes present in the network (Hage and Harary, 1995). Theonine (Thr) enzymes play important roles in the stress response to abiotic factors such as salinity, cold and drought (Rudrabhatla and Rajasekharan, 2002; Diédhiou et al., 2008), in addition to the different processes related to plant growth, such as cell division and the regulation of several phytohormones (Rudrabhatla and Rajasekharan, 2002) and carbon flux (Zeh et al., 2001).

In this work, we identified many genes involved in the response to drought, showing the importance of this element for the development of rubber trees, as already reported by Conforto (2008). Conson et al. (2018) and Souza et al. (2019) showed that the environments in which the populations used in this work are grown are environments with constant water deficit, which was expected because they are escape areas, which presents different climate of their natural habitat but where the rubber tree has adapted well . In the context of climate change, the discovery of genes involved in responding to water stress is of great value since forecasts show that in the near future areas suitable for planting today may become unsuitable (Ray et al., 2016). Most likely, these changes will occur mainly in the water regime, which can lead to the death of many woody plants (Adams et al., 2009).

Despite the limitations of the GWAS in identifying genes related to quantitative traits, the multiomics strategy employed in this study allowed us to explore the main genes that putatively define this phenotype from a holistic perspective, expanding this investigation and supplying a large reservoir of data. Using the integration of GWAS with coexpression networks and enzyme networks, we were able to elucidate the main relationships of these major genes and their products in a more complete way, mainly considering the limitations of GWAS in the identification of regions of QTLs with small effects. With the functional modules defined, we can gain insight into the genes that work together. In addition to the understanding that the definition of SD is based on the interaction of several processes, we have identified 6 functional modules. Even with more than one process, all these interactions work together, as we can see in the network shown in Figure 6. In addition, we can see the robustness of these results, which show correlations with previously published QTL maps (Conson et al., 2018). Posttranslational inferences were made regarding the relationships identified in the enzymatic network, which allowed us to identify new and important gene products that were previously unidentified. All these results show the importance of these integrative studies that correct the limitations of each individual technique.

This work is the first initiative that integrates multiomics in the study of QTLs in *H. brasiliensis*. Using this approach, we were able to access all important molecular levels for the definition of SD. Despite the great economic importance of the species, as it is the only one capable of producing natural rubber in sufficient quantity and quality to supply the world market for this product (Ding et al., 2020), its genetic studies are still quite limited due to the complexity of its genome (Tang et al., 2016), its great genetic variability (De Souza et al., 2018) and the large areas needed for its plantation. Despite all these limitations, this work overcomes these difficulties, producing data, results and new methodological perspectives for future genomic studies in this species and identifying markers and genes useful for genetic improvement.

## Conflict of Interest

The authors declare that the research was conducted in the absence of any commercial or financial relationships that could be construed as a potential conflict of interest.

## Author Contributions

FF and AA performed all the analyses and wrote the manuscript; CC assisted in the genotypic data analyses; PG, VG and EJ conducted the field experiments; and AS, LS and RF-N conceived the project. All authors reviewed, read and approved the manuscript.

## Funding

The authors gratefully acknowledge the Fundação de Amparo à Pesquisa do Estado de São Paulo (FAPESP) for Ph.D. fellowship to FF (18/18985-7) and AA (2019/03232-6); The Coordenação de Aperfeiçoamento do Pessoal de Nível Superior (CAPES) for financial support (Computational Biology Program and CAPES-Agropolis Program); and the Conselho Nacional de Desenvolvimento Científico e Tecnológico (CNPq) for research fellowships to AS and PG.

## Abbreviations

ABA: Hormone abscisic acid
BLUP: Best linear unbiased predictor CINV2: Alkaline/neutral invertase CINV2 CPM: Counts per million
DEGs: Differentially expressed genes EC: Enzyme commission
EST: Expressed sequence tag
FarmCPU: Fixed and random model Circulating Probability Unification FBX5: Arabidillo 1 protein
GBS: Genotyping-by-sequencing GO: Gene Ontology
GOLS2: Protein galactinol synthase 2 GSO1
GSO1: LRR receptor-like serine/threonine-protein kinase GWAS: Genome-wide association studies
H²: Broad heritability
HGN1: Protein glucan endo-1,3-beta-glucosidase INV: Invertase enzyme
JUB1: Transcription factor jumgbrunnen 14 KEGG: Kyoto encyclopedia of genes and genomes kNNI: k-nearest neighbor imputation
LD: Linkage disequilibrium
LRT: Likelihood ratio test
MAF: Minimum allele frequency MAS: Marker-assisted selection
MIOX: Substrate of the myo-inositol oxygenase ML4: MEI2-like 4
NGS: Next-generation sequencing PCA: Principal component analysis PE: Paired-end
PUB23: Protein E3 ubiquitin-protein ligase PUB23 Q-Q plot: Quantile-quantile graph
QTLS: Quantitative trait loci
R²: Squared Pearson correlation
RAPTOR1: Regulatory-associated protein of TOR 1 RE1 RNA-seq: RNA-sequencing
RE1: Retrovirus-related Pol polyprotein from transposon SD: Stem diameter
SNPs: Single nucleotide polymorphism snpsGWAS: SNPs discovered by GWAS snpsLD: GWAS LD-associated markers SSRs: Microsatellite markers
SUS: Sucrose synthase enzyme TE: Transposons elements
Thr: Theonine
thrC: Threonine synthase
TOM: Topological overlap measure
UPGMA: Unweighted Pair-Group Method using Arithmetic Avarages USP: UDP-sugar pyrophosphorylase
WGCNA: Weighted gene correlation analysis

## Notes

### Competing Interest Statement

The authors have declared no competing interest.

