## Supplementary figures for "Unravelling Rubber Tree Growth by Integrating GWAS and Biological Network-Based Approaches"

### *Supplementary Material*

#### 1 Supplementary Figures

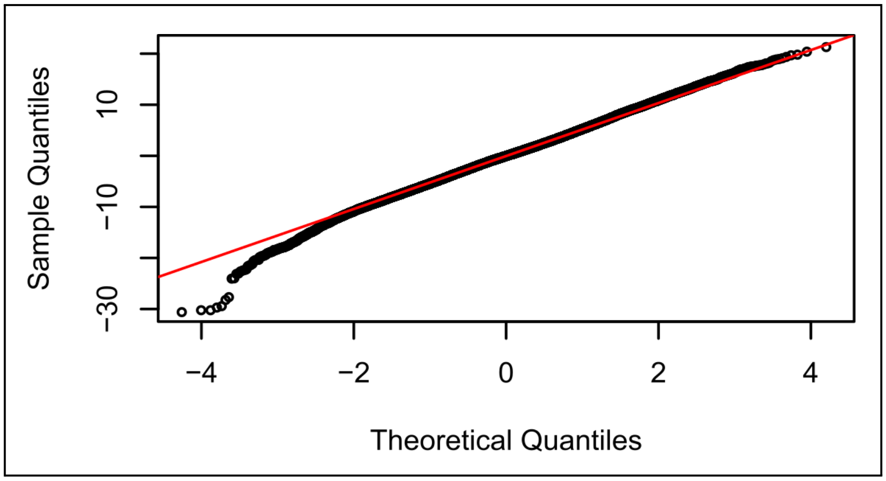


**Supplementary Figure 1**. Quantile-Quantile (QQ) plot showing the normality of the residuals of the mixed models used for the analysis of the phenotypic data.

**Supplementary Material**

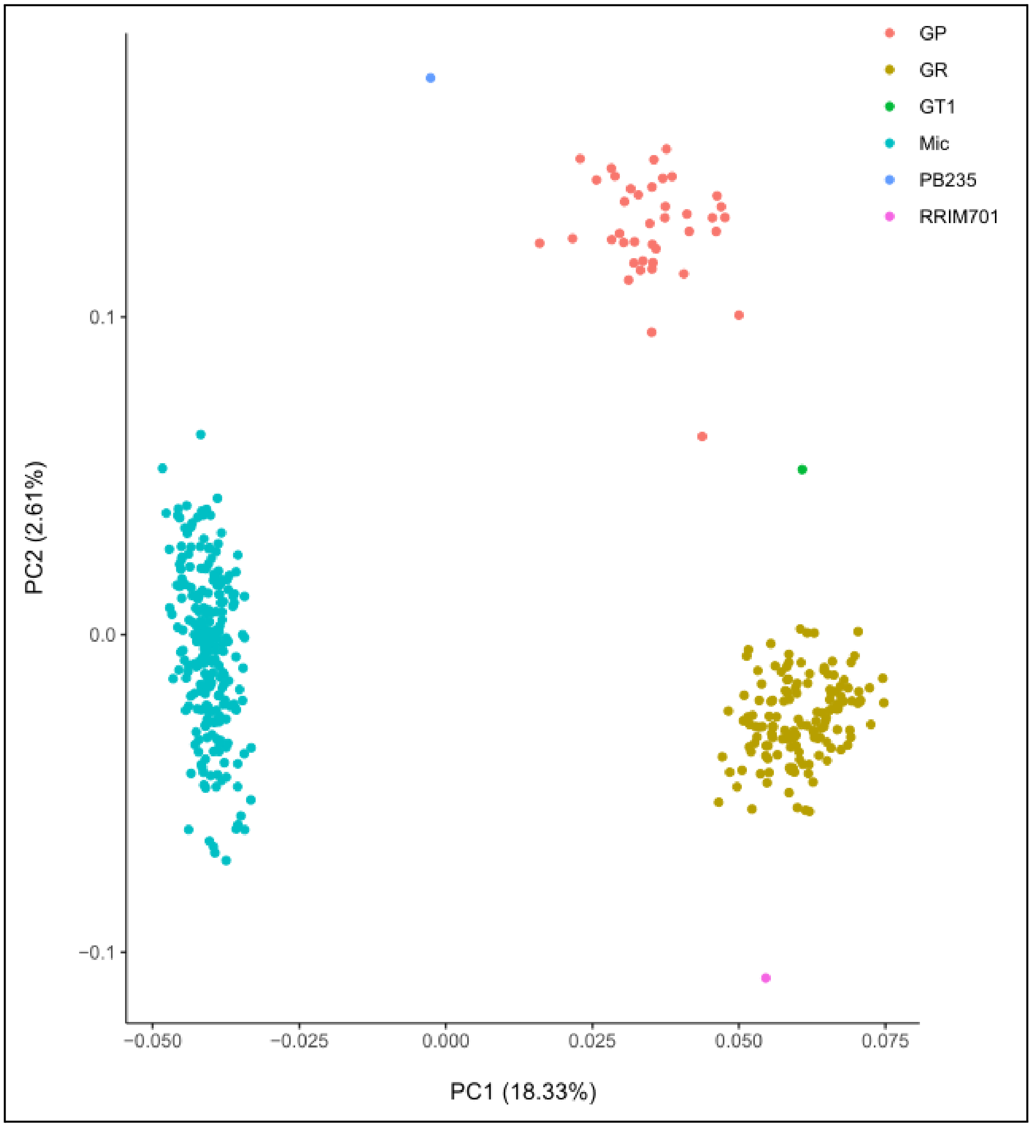


**Supplementary Figure 2**. Principal component analysis (PCA) scatter plot with the genomic data represented in two principal components (PCs). Each point represents a genotype from the populations (i) GP (GT1xPB235); (ii) GR (GT1xRRIM701); and (iii) Mic (PR255xPB217); and from the genotypes GT1, PB235 and RRIM701.

**Supplementary Material**

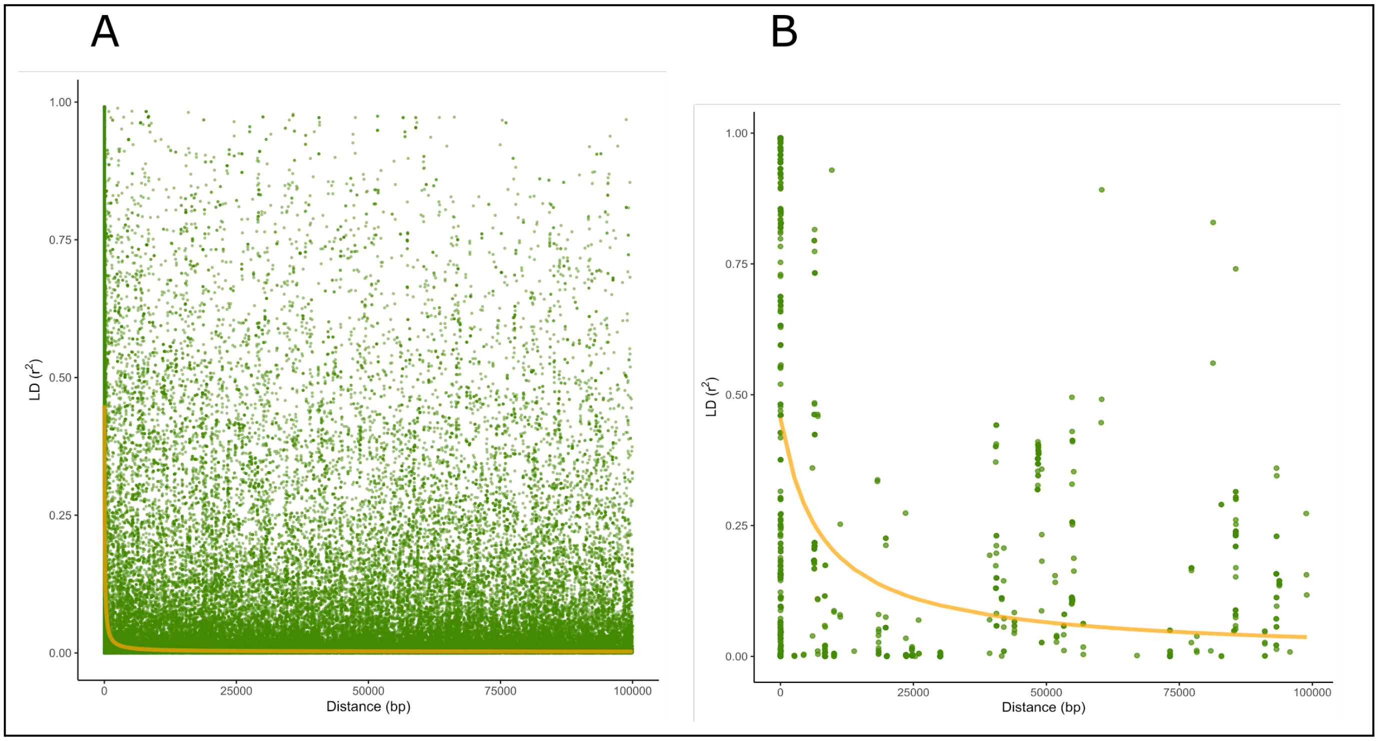


**Supplementary Figure 3**. (A) Linkage disequilibrium (LD) decay along the entire genome; (B) LD decay only in regions containing transposable elements.

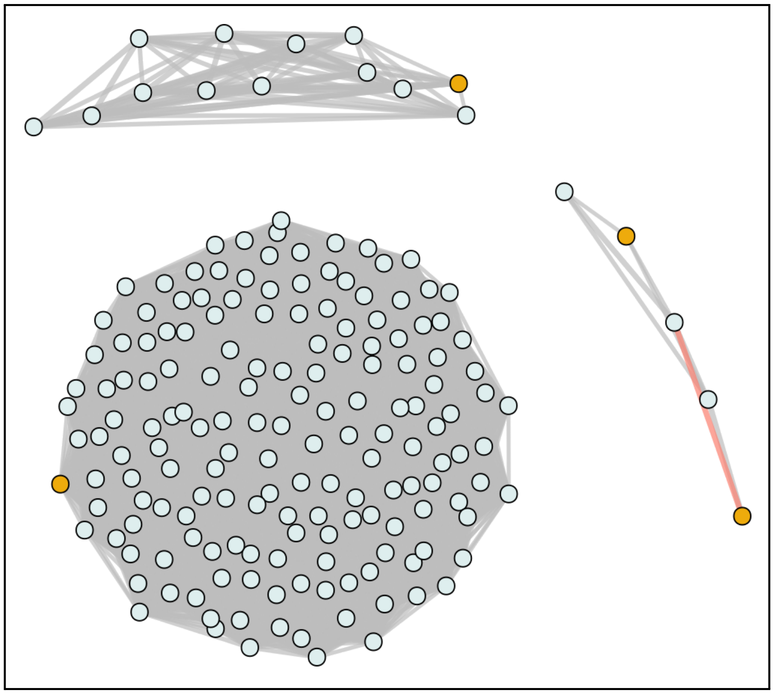

**Supplementary Figure 4**. Correlation network between the SNPs identified by the GWAS, in yellow, and all other SNPs in grey. Edges indicate a correlation above 70%, being colored in red when there is a negative correlation.

**Supplementary Material**

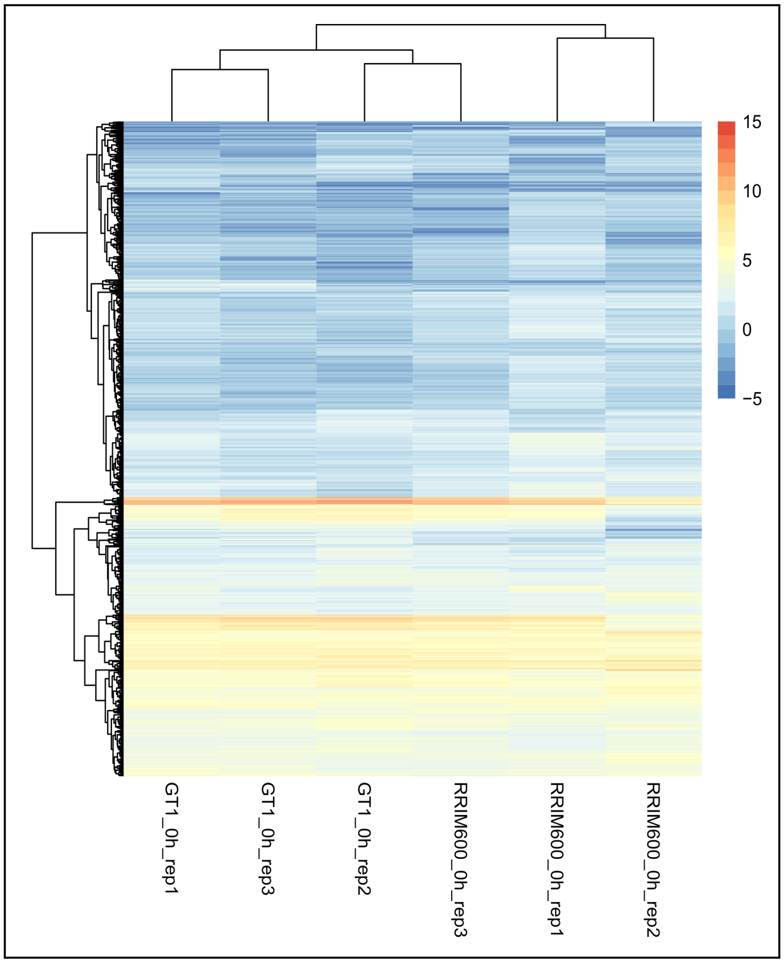


**Supplementary Figure 5**. Heatmap of the expression of genes identified in the gene coexpression network modules selected. The quantification is presented in logarithmic scale.

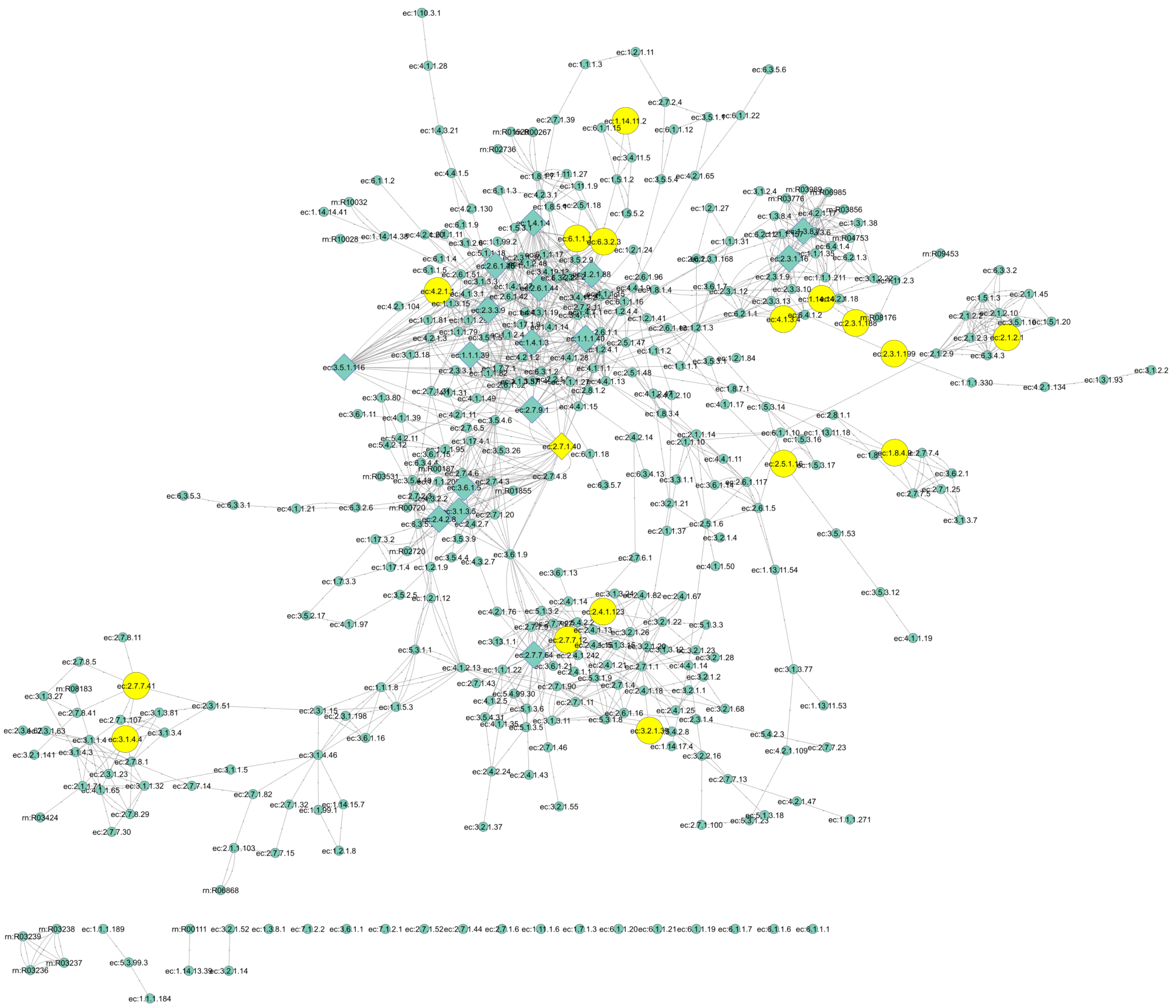


**Supplementary Figure 6**. Enzyme network with labels shown. The yellow nodes represent the enzymes discovered in the coexpression modules, and the rectangular nodes indicate the enzymes with the highest centrality values.

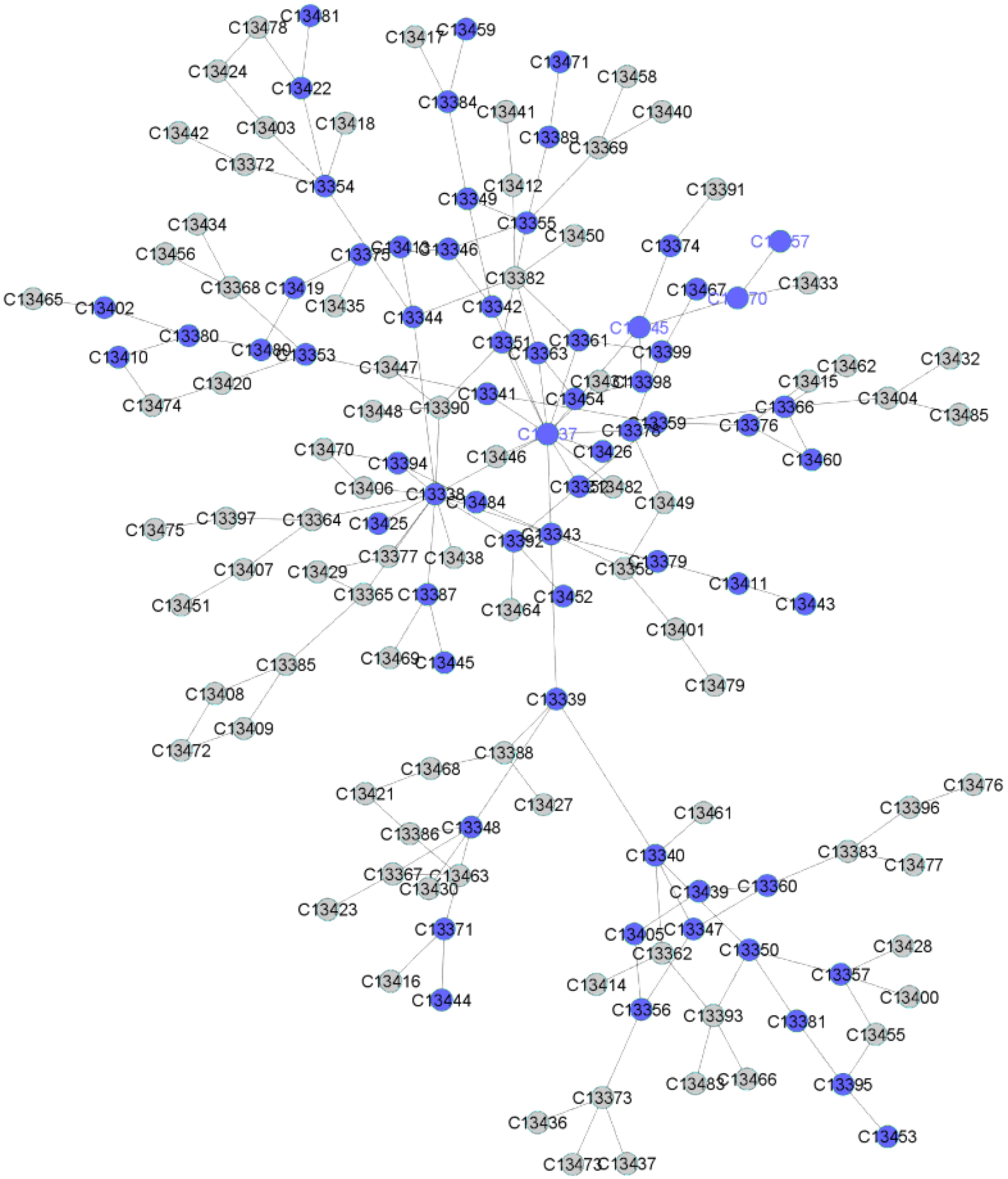


**Supplementary Figure 7**. Enzyme network with labels shown (communities). The blue nodes are represented by communities containing enzymes discovered in the coexpression modules.
